# A numbers game: Mosquito-based arbovirus surveillance in two distinct geographic regions of Latin America

**DOI:** 10.1101/2024.03.15.585246

**Authors:** Jacqueline Mojica, Valentina Arévalo, Jose G. Juarez, Ximena Galarza, Karla Gonzalez, Andrés Carrazco, Harold Suazo, Eva Harris, Josefina Coloma, Patricio Ponce, Angel Balmaseda, Varsovia Cevallos

## Abstract

*Aedes* mosquitoes, as vectors of medically important arthropod-borne viruses (arboviruses), constitute a major public health threat that requires entomological and epidemiological surveillance to guide vector control programs to prevent and reduce disease transmission. In this study, we present the collaborative effort of one year of mosquito-based arbovirus surveillance in two geographically distinct regions of Latin America (Nicaragua and Ecuador). Adult female mosquitoes were collected using backpack aspirators in over 2,800 randomly selected households (Nicaragua, Ecuador) and 100 key sites (Nicaragua) from eight distinct communities (Nicaragua: 2, Ecuador: 6). A total of 1,358 mosquito female pools were processed for RNA extraction and viral RNA detection using real-time RT-PCR. Ten positive dengue virus (DENV) pools were detected (3 in Nicaragua and 7 in Ecuador), all of which were found during the rainy season and matched the serotypes found in humans (Nicaragua: DENV-1 and DENV-4; Ecuador: DENV-2). Infection rates ranged from 1.13 to 23.13, with the Nicaraguan communities having the lowest infection rates. Our results demonstrate the feasibility of detecting DENV-infected *Aedes* mosquitoes in low-resource settings and underscore the need for targeted mosquito arbovirus sampling and testing, providing valuable insights for future surveillance programs in the Latin American region.

## Introduction

*Aedes aegypti* is the main vector of medically important arthropod-borne viruses (arboviruses), such as dengue, chikungunya and Zika viruses. As *Aedes* mosquitoes continue to expand their global distribution (Messina et al., 2019), almost four billion people are now at risk for viral infection (WHO, 2023). This constitutes a major public health threat that requires entomological and epidemiological surveillance to guide vector control programs to prevent and reduce burden of disease. However, detecting early signs of arbovirus transmission is challenging, and mosquito-based arbovirus surveillance (*i*.*e*., detecting arboviruses in mosquitoes) is viewed as a tool that might provide early warning of impending outbreaks (Grubaugh et al., 2015). Achieving early detection of arboviruses in resource-limited settings presents challenges for sustainability since it requires the use of nucleic acid detection technology (real time RT-PCR or quantitative RT-PCR [qRT-PCR]) (Ramírez et al., 2018), which can be an expensive endeavor. Nonetheless, mosquito-borne arbovirus surveillance might provide a finer level of detail regarding vector transmission in local settings and help guide public health policy related to vector control tools. In this study, we present the results of one year of mosquito-based arbovirus surveillance in two distinct geographical regions of Latin America and our experience with implementation in resource limited-settings. This project was carried out as part of the Asian-American Centers for Arbovirus Research and Enhanced Surveillance (A2CARES) of the NIH Centers for Research on Emerging Infectious Diseases (CREID) network.

## Methods

Study sites and design. This collaborative study was conducted in two geographically distinct regions; namely, Nicaragua (Central America) (Figure A1.1) and Ecuador (South America) (Figure A1.2). In Nicaragua, we worked in two districts within the capital city of Managua: District 2 (classified as urban) and District 3 (classified as urban-rural). The study focused on the catchment area of one public sector health center (District 2) and two health posts (District 3). Together, these districts comprise over 340,000 inhabitants and an overall population density of 3,800 people/km^2^ (INIDE, 2021). In Ecuador, our research focused on six communities (Borbón (urban-rural), Colon Eloy, Santa Maria, Santo Domingo, Maldonado and Timbiré (rural) located in the northwestern province of Esmeraldas. Based on our study census, the population in the study communities is ∼20,000 individuals. These communities have a combined population density of 41 people/km^2^ (Zambrano, 2023). Both districts in Nicaragua and the communities in Ecuador encompassed low-income neighborhoods, with varying degrees of urban and rural characteristics, as well as differential access to healthcare services. The overall climate in both regions is tropical with marked rainy seasons (Nicaragua: June-December; Ecuador: March-July) (Thorsen, 2015) that overlaps with the dengue transmission cycle.

The entomological surveillance programs established for both sites are part of ongoing community-based cohort studies evaluating arboviruses in each region. In Nicaragua, we randomly selected a subset of 500 households from each district (of 2,166 and 1,000 households in the parent cohort study of District 2 and 3, respectively) and ∼50 key sites (*i*.*e*., tire shops, cemeteries, and schools) from each district. This selection was done using a framework for patches of risk based on landscape features. In Ecuador, an average of 30% of all community households were randomly selected. The number of households varied from 356 to 2,880 depending on the community, for a total of 546 households surveyed.

### Adult *Aedes aegypt*i surveillance

Adult mosquito collections were performed from February 2021 to December 2022. In Nicaragua, each household and key site was visited twice, once during the dry season (February-March) and once during the rainy season (September-December) of 2022. In Ecuador, each household was visited 17 times, with monthly surveys from February 2021 to July 2022, except for June 2022 due to a flooding event. In both sites, we used backpack aspirators (Prokopack), with slightly distinct collection methodologies. In Nicaragua, each household and key site was surveyed counterclockwise searching all available spaces (both indoors and outdoors), with field activities from 8 am to 5 pm. In Ecuador, households were surveyed using a 10-minute indoor procedure that prioritized areas of human presence (*e*.*g*., bedrooms, kitchens, bathrooms, living rooms, etc.), with field activities from 8 am to noon.

### Sample processing and laboratory procedures

At both research sites, all adult mosquitoes were transported in coolers to our field facilities. Mosquito identification and sorting were performed by highly trained field entomologists. All *Aedes aegypti* mosquitoes were sorted by trapping location (indoor and outdoor), sex (males and females), and feeding status (non-fed and bloodfed). In Nicaragua, female mosquitoes were grouped into pools of ≤20/mosquitoes by household. In Ecuador, pools consisted of mosquitoes from a group of 15 households, with each pool containing ≤30 mosquitoes. For both sites, each pool was preserved in RNA*later* (ThermoFisher) and stored at -80°C until sample processing for molecular testing of arboviruses.

*Aedes aegypti* pools were initially macerated using liquid nitrogen and resuspended in 300µL of PBS 1X and Triton X-100 (in Ecuador and Nicaragua), prior RNA extraction. Pools were processed for RNA extraction using QIAamp Viral RNA kits and RNeasy Mini Kit (Qiagen, Germany) following the manufacturer’s specifications. Amplification of genetic material to detect dengue (DENV), zika (ZIKV) and chikungunya (CHIKV) arboviruses was performed using ZDC Multiplex RT-PCR Assay Kit (Bio-Rad, USA) (Waggoner et al., 2016). Afterwards, Real Time RT-PCR Assay was used to detect DENV serotype (DENV-1 to DENV-4) in the DENV positive pools (Santiago et al., 2013; Waggoner et al., 2013).

### Infection rate analysis

The infection rates of mosquito pools provide an estimation of the probability of individual mosquito positivity, including point (P) and interval estimations (95% Confidence Interval [CI]). The infection rate was calculated only for the collection period during the rainy season for both sites (Nicaragua: September–December; Ecuador: February–July 2022). To account for pool size variability, we used the Firth estimator per 1,000 females (Hepworth and Biggerstaff, 2017) using the R package PooledInfRate v1.5.

## Results

### Adult collections

In total, we collected 12,162 (Nicaragua: 3,996 [household: 943, key site: 3,053]; Ecuador: 8,154) adult *Ae. aegypti* specimens, of which 6,244 were females (Nicaragua: 1,878; Ecuador: 4,366) with 2,200 bloodfed females (Nicaragua: 94; Ecuador: 2,106) (Table 1). During the rainy season period, a total of 2,624 female *Ae. aegypti* were collected (Nicaragua: 1823; Ecuador: 801). We observed a difference in the average number of captured females between sites and trapping locations. In Nicaragua, indoor collections averaged 0.29 female *Ae. aegypti* per household, while in Ecuador we obtained a higher average of 1.47 female *Ae. aegypti* per household. Outdoor collections in Nicaragua showed the lowest value, with an average of 0.13 female *Ae. aegypti* per household. Key site surveillance in Nicaragua was particularly productive, with 1,450 female *Ae. aegypti* mosquitoes, representing 75% of all female collections in Nicaragua during this season. Notably, in the cemetery of District 2, where 794 tombs were inspected, we collected over 40% (1,216) of all female *Ae. aegypti* in Nicaragua.

**Table 1.** Female *Aedes aegypti* adult collection in Nicaragua and Ecuador.

|  | Nicaragua | Ecuador |
| --- | --- | --- |
| **Household<sup>1</sup>** |  |  |
| Indoor | 296 (0.3) | 4,366 (1.5) |
| Outdoor | 132 (0.1) | - |
| **Key sites** |  |  |
| Indoor | 132 | - |
| Outdoor | 1318 | - |
| **Total females** | 1,878 | 4,366 |
| Un-engorged | 1,784 | 2,260 |
| Bloodfed | 94 | 2,106 |
| **Infection rate<sup>2</sup>** |  |  |
| Urban<sup>3</sup> | 1.31 (95% CI = 0.2–4.3) |  |
| Urban-Rural<sup>4</sup> | 3.43 (95% CI = 0.2-16.4) | 7.38 (95% CI = 2.8-16.3) |
| Rural<sup>5</sup> |  | 9.06 (95% CI = 0.5-45.1)<sup>6</sup><br>23.13 (95% CI = 1.4-115-9)<sup>7</sup> |

<sup>1</sup>Average female per household in ( ).
<sup>2</sup>Data calculated only for the rainy season collection period of 2022 (Nicaragua: September–December; Ecuador: February–July).
<sup>3</sup>Urban: Nicaragua (District 2)
<sup>4</sup>Urban-Rural: Nicaragua (District 3)– Ecuador (Borbón)
<sup>5</sup>Rural: Ecuador (<sup>6</sup>Maldonado; <sup>7</sup>Timbire)

### Mosquito-based arbovirus surveillance

A total of 1,358 pools (Nicaragua: 871; Ecuador: 487) were tested during the entire study period. Mosquito positivity was only observed during the rainy season (DENV transmission season), with 10 positive pools for arboviruses (all DENV) across sites and the earliest detection two months after the start of the rainy season. In Nicaragua, during the rainy season we processed 824 pools (1809 females) with 3 DENV-positive pools. Two positive pools were found in District 2: one was collected in a household (DENV-1) and the other in a key site (cemetery, DENV serotype not identified) (Figure 1B). A single positive female was identified in District 3 with a co-infection of DENV-1 and DENV-4, collected in a household (Figure 1C). It is worth noting that all positive mosquitoes were visibly non-engorged “non-fed” females. In Ecuador, during the 2022 rainy season we processed 64 pools (1072 females; 576 bloodfed) with 7 DENV-positive pools all of which were DENV-2. Five positive pools were found in the community of Borbón (Figure 1D), one in Timbiré (Figure 1E), and one in Maldonado (Figure 1F).

**Figure 1.**
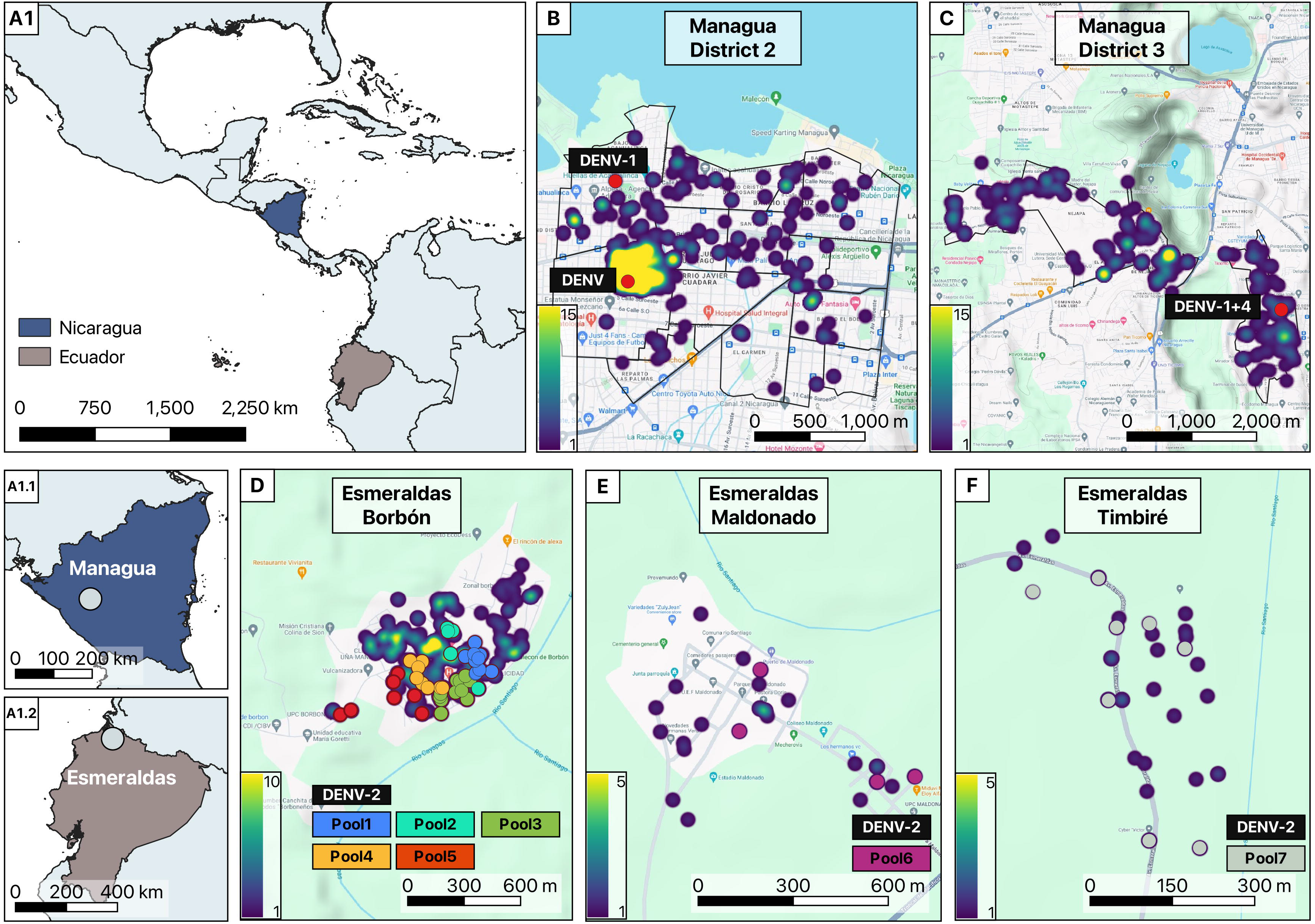
Geographic location of the research sites in A1.1) Managua, Nicaragua, and A1.2) Esmeraldas, Ecuador, with positive arbovirus detection in female *Ae. aegypti*. Gradient pattern color shows a heatmap of *Ae. aegypti* abundance by research site. B) District 2 of Managua, with positive detection for DENV and DENV-1. C) District 3 of Managua, with positive detection of a single co-infection of DENV-1 and DENV-4. D) Commercial area of Borbón in Esmeraldas, with positive detection of DENV-2. E) Town of Maldonado in Esmeraldas, with positive detection of DENV-2. F) Town of Timbiré in Esmeraldas, with positive detection for DENV-2. Nicaragua: Red dots indicated positive pools per household. Ecuador: different color dots represent different pools. Maps were generated using QGIS v3.34 with publicly available administrative boundaries and satellite imagery from OpenStreetMaps (OpenStreetMap contributors, 2023).

Mosquito infection rates showed a clear difference between research sites and communities. In Nicaragua, the infection rate in District 3 was P = 3.43 (95% CI, 0.2–16.4). In District 2, infection rate was slightly lower, with P = 1.31 (95% CI, 0.2–4.3). In contrast, Ecuador exhibited notably higher infectious rates, with Borbón having P = 7.38 (95% CI, 2.7–16.3). In Maldonado, the infection rate was P = 9.05 (95% CI, 0.5–45.2). In Timbiré, the infection rate was P = 23.13 (95% CI, 1.4–115.9). These variations underscore the heterogeneity of arbovirus transmission dynamics observed in different regions and communities within our study.

## Discussion

Mosquito-based arbovirus surveillance is viewed as a tool that can help guide vector control programs to prevent virus outbreaks (Leandro et al., 2022). Nevertheless, in resource-limited settings, its implementation can impose a significant operational burden if not applied appropriately. We conducted a thorough evaluation of mosquito-based arbovirus surveillance within two distinct geographical settings in Latin America with varying degrees of urbanization and accessibility. Our results show that mosquito-based arbovirus surveillance employing random household sampling might not be a sustainable methodology for arbovirus detection transmission and suggest the need to re-evaluate mosquito-based arbovirus surveillance approaches to ensure the most effective use of resources in combatting arbovirus transmission. Entomological surveillance efforts in over 2,800 households and over 100 key sites yielded only 10 positive pools using a random sampling approach. No tested pool showed positivity outside of the rainy season. Our findings highlight the importance of establishing a defined surveillance timeframe or targeted sampling approach in hotspots, in which active human dengue cases are identified, which others have shown increases mosquito positivity rates (Krokovsky et al., 2022). As such, implementing arbovirus testing in mosquitoes should be done using a more targeted approach that could be more cost-effective. However, sampling around hotspots of cases would no longer be predictive of emerging outbreaks. Additionally, the use of mass pooling at the community level (Tang et al., 2020) and targeted sampling based on human movement patterns (Stoddard et al., 2009) or mosquito dispersal (Juarez et al., 2020) could improve mosquito-based surveillance tools. It has also been observed that even across close distances, viral loads in *Ae. aegypti* populations can vary widely and impact infection and vector competence rates (Godoy et al., 2021). The approaches, mentioned above, could not only reduce operational costs but also guide vector control activities over a broader area. A clear limitation of our study is that our entomological protocols were not identical, reducing the comparability between research sites. However, we show that despite applying two different approaches, we did not obtain better outcomes with a particular approach.

Importantly, if mosquito-based arbovirus surveillance is to be conducted, our results show that this procedure needs to be expanded beyond the household to other mosquito key sites where mosquitoes abound or where people congregate, in order to increase the odds of detecting infected specimens. The ability to address the challenging task of detecting infected *Ae. aegypti* mosquitoes, which can be influenced by numerous variables including low viral loads and assay limit of detection (Kirstein et al., 2021), requires novel cost-effective tools that allow for early viral detection on a fine scale. We hope that our results provide evidence to support the importance of targeted mosquito arbovirus sampling and testing. These findings can provide valuable insights for future surveillance programs in the Latin American region. By adopting a more strategic and efficient approach, such as hot spot area surveillance from previous years, it should be possible to enhance the capacity of vector control programs to detect and respond to arbovirus outbreaks, contributing to improved disease control and prevention efforts.

## Acknowledgments

We would like to thank all the members of our A2CARES study teams in Nicaragua and Ecuador who were involved in the collection of entomological samples and laboratory procedures. We thank the residents who allowed entomological collections in and around their homes.

## Data availability

See Supplemental Dataset and https://github.com/jgjuarez/Mojica_Entovirology/tree/main for Rcode.

## Author Contributions

**JM:** Methodology; Investigation; Data curation; Formal analysis; Writing – original draft. **VA:** Methodology; Investigation; Data curation; Writing – review & editing. **JGJ:** Conceptualization; Data curation; Formal analysis; Writing – original draft. **XG:** Methodology; Investigation; Data curation; Formal analysis; Writing – original draft. **KG:** Methodology; Investigation. **AC:** Methodology; Investigation. **HS:** Conceptualization; Investigation. **VA:** Methodology; Investigation. **EH:** Conceptualization; Methodology; Funding acquisition; Writing – review & editing. **JC:** Conceptualization; Methodology; Funding acquisition; Writing – review & editing. PP: Conceptualization; Methodology; Data curation. **AB:** Conceptualization; Methodology. **VC:** Conceptualization; Methodology; Writing – review & editing.

## Ethics

Protocols for the collection and testing of samples were reviewed and approved by the Institutional Review Boards (IRB) of the University of California, Berkeley (PDCS: 2010-09-2245; A2CARES: 2021-03-14191) and the Nicaraguan Ministry of Health (PDCS: CIRE 09/03/07/-008.Ver25; A2CARES: CIRE 02/08/21-114 Ver.4) and Universidad San Francisco de Quito (2017-0159M).

